# 5-HT_4_ receptor ligand RS67333 modulates striatal acetylcholine and dopamine release via inhibition of acetylcholinesterase

**DOI:** 10.64898/2026.06.24.733606

**Authors:** Qinbo Qiao, Wenhui Wu, Stephanie J. Cragg

## Abstract

Serotonin 5-HT4 receptors (5-HT_4_Rs) have emerged as potential therapeutic targets in neuropsychiatric and neurodegenerative disorders by modulating circuits that shape mood, cognition, and motor function. Ligands for 5-HT_4_Rs can modify dopamine (DA) and acetylcholine (ACh) transmission but mechanisms and circuits have not been fully resolved. Some 5-HT_4_R agonists have been suggested to have effects that include inhibition of acetylcholinesterase (AChE), raising speculation that 5-HT_4_R ligands might modulate ACh and/or DA through this action. Here, we investigated the impact of RS67333, a partial 5-HT_4_R agonist, on DA and ACh release dynamics in the striatum detected *ex vivo* in mouse brain slices using fast-scan cyclic voltammetry and genetically encoded ACh sensor GRAB_ACh3.0_ respectively. We found that RS67333 significantly modulated electrically evoked DA release in dorsolateral striatum and nucleus accumbens core, effects that were abolished by a nicotinic receptor (nAChR) antagonist. In parallel, RS67333 altered evoked ACh signals by extending extracellular ACh lifetime, and correspondingly, RS67333 was found to inhibit striatal AChE enzymatic activity. By contrast, BIMU8, an alternative 5-HT_4_R ligand that did not inhibit striatal AChE, had no effect on evoked striatal ACh or DA release. These findings indicate that RS67333 modulates striatal ACh transmission, which shapes downstream regulation of DA release by nAChRs, not through 5-HT_4_Rs but through AChE inhibition. These findings emphasize the caution due in attributing functions to 5-HT_4_Rs, but also highlight an alternative pharmacological profile of some purported 5-HT_4_R ligands as AChE inhibitors of potential utility for treating ACh/DA disorders.

## Introduction

Serotonin (5-HT) type 4 receptors (5-HT_4_Rs) are G_s_/G_olf_-coupled receptors that modulate mood, cognition, and motor function (1–3) due to their expression in brain regions such as hippocampus and basal ganglia (4–6). The relatively advanced development, and availability, of 5-HT_4_R ligands makes them attractive tools for potentially treating diverse neuropsychiatric, cognitive and psychomotor disturbances (2,3,7–13). For example, RS67333, a 5-HT_4_R partial agonist, enhances memory performance in object recognition tasks and ameliorates stress-induced affective behaviours in rats (7,14), while in mouse models of Parkinson’s disease (PD), 5-HT_4_R agonists RS67333 and prucalopride are reported to reduce L-DOPA-induced dyskinesias (3). 5-HT_4_R ligands have been suggested to be effective by modulating the neurotransmitters underpinning cognitive and motor control such as dopamine (DA) and acetylcholine (ACh). *In vivo* and *ex vivo* studies have shown for example that 5-HT_4_R agonists, including renzapride and (S)-zacopride, can increase extracellular DA levels in rat striatum (15–17), and that systemic or intranigral 5-HT_4_R antagonists attenuate psychostimulant-evoked DA release (18,19). However, the circuits and mechanisms involved have not been resolved. While 5-HT_4_R ligand-binding sites and gene expression (*htr4* gene) are reported in rodent and primate striatum (6,20), there is a paucity of evidence for 5-HT_4_R gene expression in midbrain DA neurons, suggesting that modulation of DA by 5-HT_4_R ligands is not direct on DA neurons, but rather is indirect via intermediary circuits (15).

In striatum, 5-HT_4_Rs have been localized rather to cholinergic interneurons (ChIs) and D_2_R-expressing GABAergic striatal projection neurons (3,6,21,22). Both ACh and GABA strongly modulate striatal DA release via actions at nicotinic ACh receptors (nAChRs) and GABA_A/B_-receptors respectively on DA axons (23,24). ChIs are in turn known to be upstream mediators for many other striatal modulators that regulate DA signaling, including opioids, glutamate, insulin, and astrocytes (25–28), making 5-HT_4_Rs on ChIs a potential locus for shaping both ACh and downstream DA signalling. However, while a broad-spectrum 5- HT receptor agonist quipazine inhibited stimulation-induced striatal ACh overflow (29), the evidence for involvement of 5-HT_4_Rs specifically is lacking. An *in vivo* microdialysis study reported that administration of selective 5-HT_4_R agonists BIMU1 and BIMU8 modulated ACh release in prefrontal cortex but not striatum (30).

Besides their efficacy at 5-HT_4_Rs, some ligands have been suggested to have a multitarget ‘pleiotropic-like’ profile, including the ability to inhibit catabolic enzyme ACh esterase (AChE). Partial 5-HT_4_R agonist RS67333 has structural similarities to AChE inhibitor donepezil and has been reported to inhibit AChE in an *in vitro assay* at submicromolar concentrations (31). The potential impact of AChE inhibition in a physiological brain setting has not yet been demonstrated, but could include significant impacts on ACh transmission, and downstream DA release, offering potential therapeutic benefits, even independently of any 5-HT_4_R efficacy. By prolonging extracellular ACh availability, AChE inhibition can rescue ACh deficits in neurodegenerative cognitive deficits in cortical/hippocampal circuits, and shape how ACh acts at striatal nicotinic ACh receptors (nAChRs) on DA axons to modulate DA release (32,33). 5-HT_4_R ligands such as exemplar RS67333 might thereby modify ACh and DA signalling through mechanisms independent of 5- HT_4_R function.

In the present study, we explored the impact of RS67333 on striatal DA and ACh release dynamics, and identified underlying mechanisms. We find that RS67333 modifies DA release in dorsal and ventral striatum, mediated via a prolongation of extracellular ACh signals acting at nAChRs on DA axons, owing to inhibition of striatal AChE. By contrast, an alternate 5-HT_4_R ligand BIMU8, lacked AChE inhibitory properties, and failed to have any effect on striatal DA or ACh release. These findings reveal that the action of RS67333 as a potent AChE inhibitor and not as a 5-HT_4_R ligand are responsible for modulation of striatal ACh and downstream DA transmission. These findings also indicate that 5-HT_4_Rs do not directly modulate DA release from DA axons. More widely, these findings suggest a re-evaluation of the potential applications of purported 5-HT_4_R drugs across the drug group, to take into account potential actions on AChE.

## RESULTS and DISCUSSION

### DA release in DLS and NAcC is inhibited by 5-HT_4_R agonist RS67333

We first investigated the impact of 5-HT_4_ receptor agonist RS67333 on evoked DA release in dorsolateral striatum (DLS) and in nucleus accumbens core (NAcC) in acute coronal slices of mouse striatum, using FCV at carbon-fibre microelectrodes to detect DA released by electrical pulses delivered as single pulses (1p) or in short trains of 5-pulses (5p) at 100 Hz. The ratio of release evoked by these two protocols (5p:1p) is useful for exposing dynamic changes in DA release probability, including those that can arise from changes to the action of ACh acting at nAChRs on DA axons (34–37). We found that RS67333 modifies DA release in DLS and NAcC. In DLS, application of RS67333 (10 µM) significantly reduced [DA]_o_ evoked by a single pulse (1p), approximately halving peak [DA]_o_ compared to pre-drug conditions (**Fig. 1A-B**; *F*_*(1,10)*_=12.60, *p=*0.0137, RM ANOVA main effect of the drug). [DA]_o_ evoked by 5p/100 Hz was not reduced significantly from pre-drug levels in DLS (**Fig. 1A-B**; *p=*0.0690, Fisher’s LSD test), and correspondingly, the ratio of peak [DA]_o_ evoked by 5p versus 1p (5p:1p ratio) was increased by RS67333 from ∼1.1 to ∼2 (**Fig. 1C**; *t*(5)=4.240, *p=*0.0082; Student’s paired *t*-test). In NAcC, RS67333 significantly reduced [DA]_o_ evoked by either 1p or 5p (100 Hz) (**Fig. 1D-E**; *F*_*(1,10)*_=18.61, *p=*0.0015, RM ANOVA main effect of the drug) and also significantly increased 5p:1p ratio from ∼1.4 to ∼2 (**Fig. 1F**; *t*(5)=4.375, *p=*0.0072; Student’s paired *t*-test).

**Figure 1.**
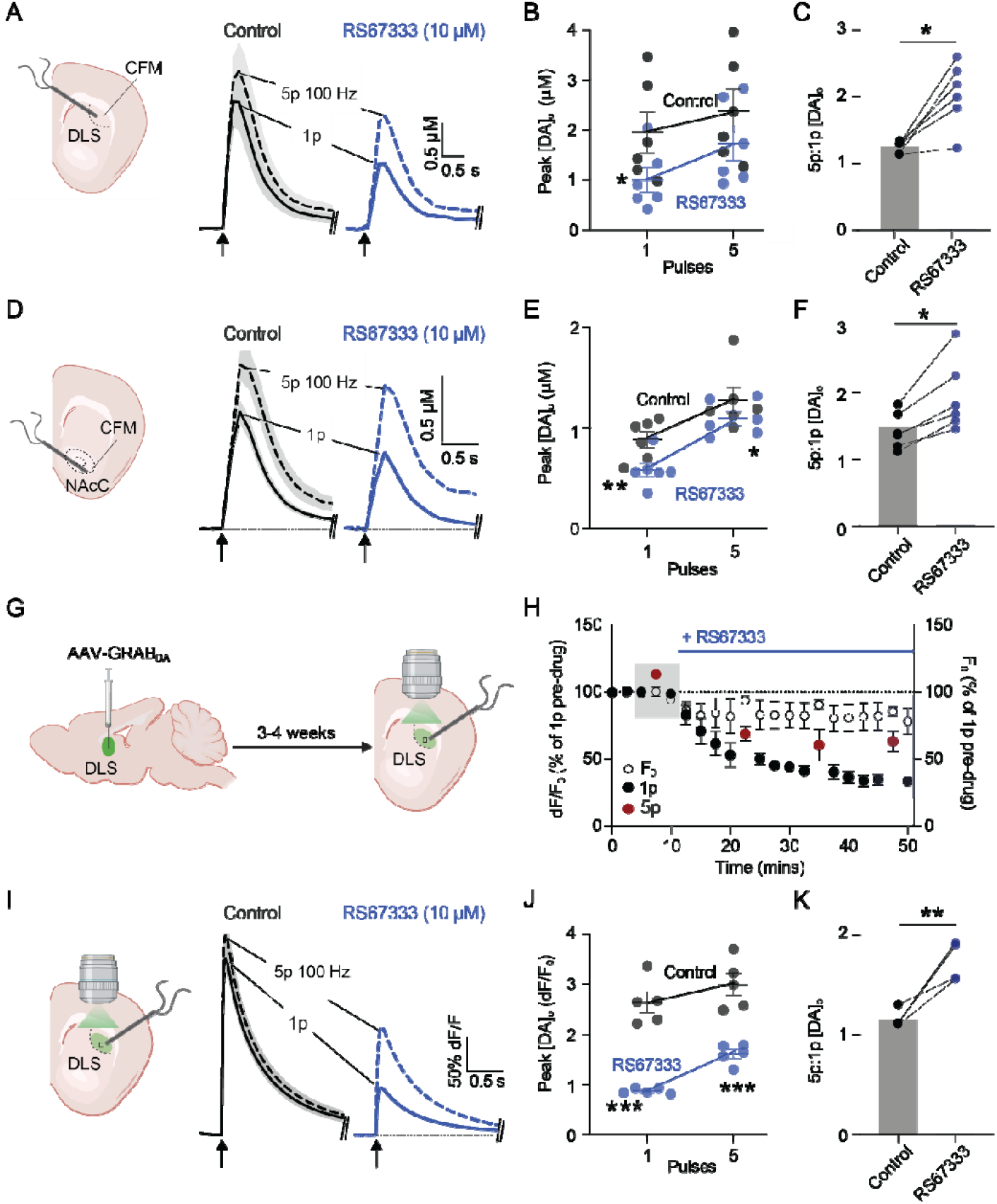
RS67333 modifies DA release and its short-term plasticity in DLS and NAcC. **(A,D)** Schematics of stimulation and recording at a carbon-fibre microelectrode (CFM) in (A) DLS or (D) NAcC, alongside mean [DA]_o_ transients ± SEM evoked (at arrows) by single pulse (1p, solid) and 5 pulses (100 Hz) (dashed), before (black) and with (blue) RS67333 (n=6, 5 mice). **(B,E)** Mean peak evoked [DA]_o_ in control (grey) or RS67333 (blue) in (B) DLS or (E) NAcC. **(C,F)** Ratios of peak [DA]_o_ released by 5p: 1p in (C) DLS or (F) NAcC. (G) Viral delivery of GRAB_DA3h_ to striatum for imaging DA release. **(H)** Normalised mean peak [DA]_o_ and baseline fluorescence (F_0_) during consecutive recordings of DA release evoked by 1p (black) and 5p (100 Hz, red), before and with RS67333, in DLS (n=5, 3 mice). **(I)** Schematic of stimulation and GRAB_DA3h_ recording, alongside mean evoked DA transients ± SEM, before (black) and with (blue) RS67333 (n=5, 3 mice) in DLS, from timepoints shaded in H. **(J)** Mean peak evoked dF/F in control (grey) or RS67333 (blue) in DLS. **(K)** Ratio of peak dF/F evoked by 5p:1p in control and drug. 2-way ANOVA, Fisher’s LSD post-hoc test, Student’s paired t-test. ^*^p < 0.05, ^**^p < 0.01, ^***^p < 0.001. Schematics created in BioRender. Cragg, S. (2026) https://BioRender.com/socz283.

To corroborate the effects of RS67333 on evoked [DA]_o_ detected with FCV, and control for changes to the sensitivity of CFMs to DA caused by the drug RS67333 (see Methods), we assessed whether RS67333 modifies evoked [DA]_o_ when detected by an alternate method, by imaging GRAB_DA_ (38), a GPCR-activation-based-DA sensor expressed after intracranial delivery of viral vector (**Fig. 1G**). In agreement with observations using FCV, RS67333 significantly reduced evoked [DA]_o_ in DLS reported by dF/F after stimulation by 1p or 5p (100 Hz), and increased 5p:1p ratio (**Fig. 1H-K**; *F*_*(1,8)*_=69.57, *p*<0.0001, RM ANOVA main effect of the drug; 5p/1p: *t*(4)=5.675, *p=*0.0048, Student’s paired *t*-test). We flag that GRAB_DA_ is a non-linear receptor-based sensor unlike the pseudo-linear detection of [DA]_o_ of FCV across this concentration range, and consequently, the magnitude of effects on evoked signals is not directly comparable for these two detection methods. More importantly, these additional observations with the GRAB-DA sensor confirm findings with FCV that RS67333 decreases DA release probability and relieves short-term depression, across striatal regions.

### Modulation of DA release by RS67333 is mediated by cholinergic interneurons in DLS and NAcC

The effects of RS6733 on DA release strongly resemble the effects of changes to nAChR activity: either a reduction in ACh action at striatal nAChRs, or an elevation of ACh that desensitizes nAChRs, reduce [DA]_o_ evoked by 1p and increase 5p:1p ratio (34,37,39). To test whether a change to nAChR activity is responsible for these effects of RS67333 on [DA]_o_, we applied a competitive nAChR antagonist, DHβE, to prevent nAChR activation, before assessing the impact of RS67333 on evoked [DA]_o_, detected with FCV. In the presence of DHβE, in DLS and NAcC, the 5p:1p ratio was larger (≥ 3) than without DHβE (compare Figs 1,2), as published previously (34). Administration of RS67333 then failed to modify [DA]_o_ evoked by 1p or 5p (100 Hz) in DLS (**Fig. 2A-B**; *F*_*(1,10)*_=1.519, *p=*0.2460, RM ANOVA main effect of the drug) or in NAcC (**Fig. 2D-E;** *F*_*(1,8)*_=0.2762, *p=*0.6134, RM ANOVA main effect of the drug). RS67333 did not modify 5p:1p ratio in either DLS (**Fig. 2C**; *t*(5)=0.1848, *p=*0.8606) or NAcC (**Fig 2F;** *t*(4)=0.3974, *p=*0.7113; Student’s paired *t*-test). These findings indicate that RS67333 modulates striatal DA release via a mechanism involving changes to ACh acting at nAChRs.

**Figure 2.**
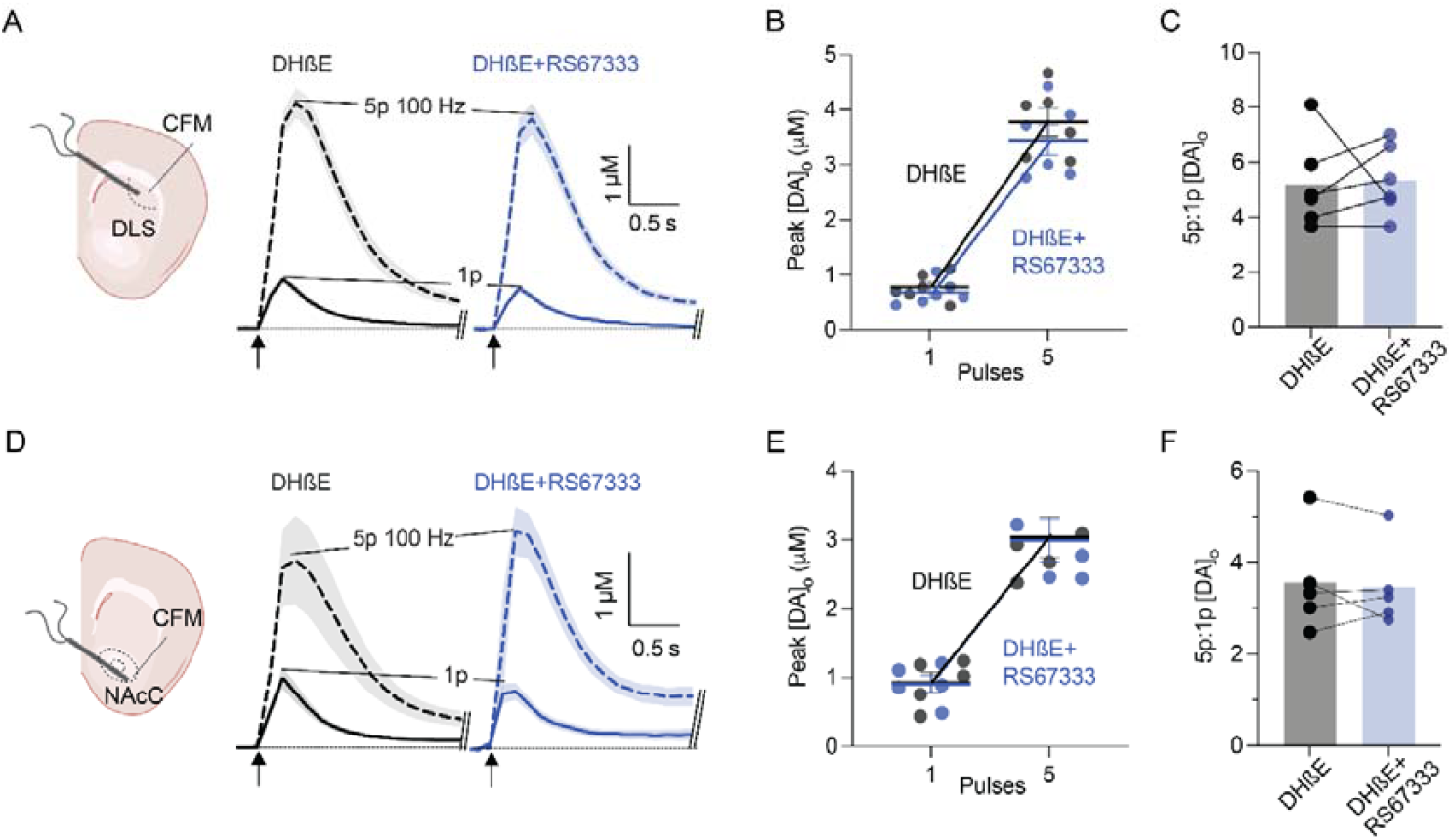
A nicotinic receptor antagonist prevents modulation of evoked striatal DA release by RS67333. **(A,D)** Schematics of stimulation and detection of evoked [DA]_o_ in (A) DLS or (D) NAcC, alongside mean [DA]_o_ transients ± SEM evoked (at arrows) by single pulse (1p, solid) and 5 pulses (100 Hz) (dashed), before (black) and with RS67333 (blue), in the presence of DHßE (n=6, 5 mice, in DLS; n=5, 4 mice in NAcC). **(B,E)** Mean peak evoked [DA]_o_ in control (grey) or with RS67333 (blue) in (B) DLS or (E) NAcC, in the presence of DHßE. **(C,F)** Ratios of peak [DA]_o_ evoked by 5p:1p in (C) DLS or (F) NAcC in the presence of DHßE. Schematics created in BioRender. Cragg, S. (2026) https://BioRender.com/socz283.

### RS67333 modifies ACh signals in DLS and NAcC as an AChE inhibitor

We investigated directly whether RS67333 modifies ACh signals. We detected extracellular ACh in mouse striatal slices by imaging fluorescence of striatally expressed GRAB_ACh3.0_ (40) (**Fig. 3A**). In DLS, RS67333 did not change peak dF/F of ACh signals evoked by 1p but slightly increased dF/F evoked by 5p (100 Hz) (**Fig. 3B-D**; *F*_*(1,8)*_=0.1285, *p*_*(1p)*_*=*0.0609, *p*_*(5p)*_*=*0.0276, RM ANOVA main effect of the drug, Fisher’s LSD test). Most notably, RS67333 significantly prolonged the extracellular lifetime of ACh signals. The integrated area under the curve (AUC) of ACh transients was tripled (**Fig. 3E-F**; *F*_*(1,8)*_*=*49.26, *p=*0.0001, RM ANOVA main effect of the drug). Single phase exponential decay curves fitted to GRAB_ACh_ signals (**Fig. 3G**) indicated that RS67333 significantly extended the ACh extracellular half-lives for either 1p or 5p stimuli (**Fig. 3H**; *F*_*(1,8)*_=7.527, *p=*0.0253, RM ANOVA main effect of the drug). A similar effect was seen in the NAcC, where RS67333 did not modify the peak df/F of ACh transients but significantly augmented the AUC and decay half-life of [ACh]_o_ (**Fig. 3I-K**; *F*_*(1,10)*_=0.3993, *p=*0.5416, RM ANOVA main effect of the drug; **Fig. 3L-M**, *F*_*(1,10)*_=15.78, *p=*0.0026, RM ANOVA main effect of the drug; **Fig. 3N, *F***_***(1,10)***_=44.33, *p*<0.0001, RM ANOVA main effect of the drug).

**Figure 3.**
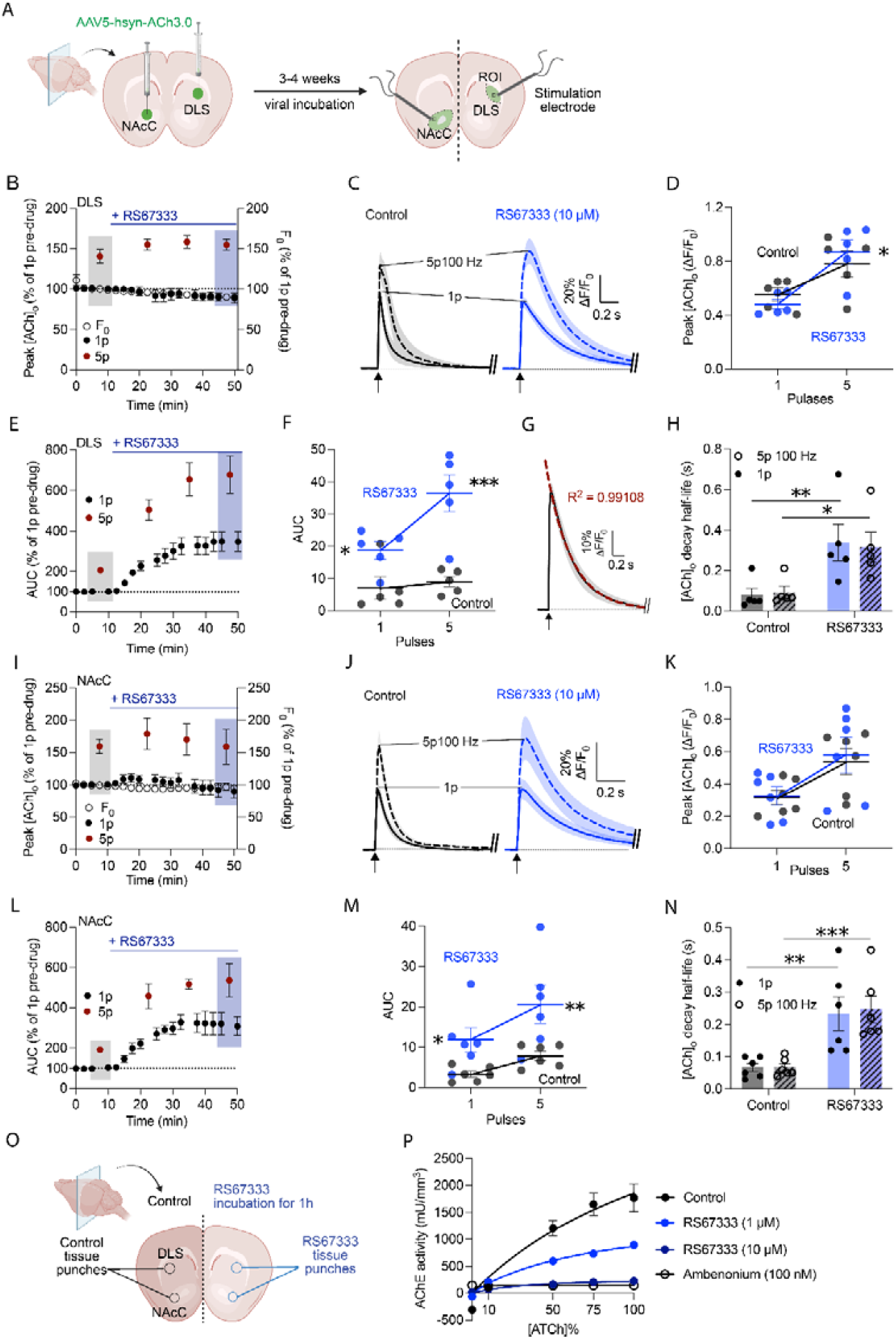
RS67333 extends extracellular ACh lifetime and inhibits AChE in DLS and NAcC. **(A)** Viral delivery of GRAB_ACh_ to striatum for imaging evoked [ACh] *ex vivo*. **(B,I)** Normalised mean peak [ACh]_o_ and baseline fluorescence (F_0_) during consecutive imaging recordings of GRAB_ACh_ signals evoked by 1p (black) and 5p (100 Hz, red), during application of RS67333 in (B) DLS or (I) NAcC (*n=*5, 3 mice). **(C,J)** Mean evoked ACh transients ± SEM, before (black) and with RS67333 (blue) (*n=*5, 3 mice) in (C) DLS or (J) NAcC, derived from the shaded timepoints in B or I. **(D,K)** Mean peak evoked dF/F in control (grey) and with RS67333 (blue) in (D) DLS or (K) NAcC. **(E,L)** Corresponding AUC plots during consecutive recordings and **(F,M)** scatter plot of AUCs values, evoked by 1p (black) and 5p (100 Hz, red), during application of RS67333 in (E,F) DLS or (L,M) NAcC (*n*=5, 3 mice). **(G)** Example of nonlinear one phase decay fitting curve (red dash line), R^2^=0.99108. **(H,N)** Decay half-lives for evoked GRAB_ACh_ signals in (H) DLS or (N) NAcC. **(O)** Schematic of tissue punches from striatal slices with or without incubation in RS67333 (1 μM or 10 µM) (*n=*4, 4 mice). **(P)** Fitted Michaelis-Menten curves of AChE activity incubated with RS67333 (1 μM and 10 μM), ambenonium (100 nM) or vehicle control. 2-way ANOVA, Fisher’s LSD post-*hoc* test, ^*^*p* < 0.05, ^**^*p* < 0.01. ^***^*p* < 0.001. Schematics created in BioRender. Cragg, S. (2026) https://BioRender.com/w6jfswd

The primary clearance mechanism for extracellular ACh is catabolism via AChE (41) and therefore we tested whether RS67333 was an inhibitor of endogenous striatal AChE, using an AChE chemiluminescent assay. RS67333 (1 μM, 10 μM) concentration-dependently inhibited AChE activity (**Fig. 3O,P**; Control: V_max_=4383 mU/mm^3^, K_m_=137.1 arbitrary units (a.u.); RS67333 (1 μM): V_max_=1538 mU/mm^3^, K_m_=76.5 a.u.; RS67333 (10 μM): V_max_=303.9 mU/mm^3^, K_m_=37.0 a.u.; Michaelis-Menten fit). The inhibition of AChE with 10 µM RS67333 was comparable to the level seen with ambenonium (**Fig. 3P**; V_max_=145.0 mU/mm^3^, K_m_=0.86 a.u.; Michaelis-Menten fit), a potent AChE inhibitor (42–44). These findings indicate that RS67333 inhibits striatal AChE, limiting degradation of released ACh and extending ACh extracellular lifetime.

Enhancements of striatal extracellular ACh action readily cause nAChR desensitization and consequent changes to downstream modulation of DA release by nAChRs across the striatum (33,45). The modulation of DA release by striatal ACh acting at nAChRs is mediated by α4α5β2- and α6β2-containing nAChRs on DA axons (46,47). These ligand-gated ion channels are activated by transient ACh release from ChIs with a range of outcomes on DA release. Brief activation of ChIs alone can be sufficient to depolarize DA axons to the threshold required for action potential generation and DA release (37), and nAChRs can also support DA release when evoked by striatal electrical stimulation (34,39,48), but they subsequently inhibit DA release driven by immediately following action potentials (36). Consequently, nAChR activation appears to support DA release probability but also short-term depression (STD) of DA release (34,36) seen as low ratios of release by multiple electrical pulses at high frequencies versus single pulses (the 5p:1p ratio identified in this study). β2-containing nAChRs also are sensitive to the duration of ACh exposure, with prolonged exposure to ACh (or exogenous agonist nicotine) leading to receptor desensitization (33,45). Thus, both a reduction in ACh levels, or sustained elevation leading to nAChR desensitization, can reduce [DA]_o_ evoked by single pulses and relieve STD to elevate [DA]_o_ evoked by pulse trains. Prior studies using AChE inhibitors such as donepezil and ambenonium have demonstrated that elevated ACh levels can desensitize nAChRs and modify striatal DA output in rodent models, both *ex vivo* and *in vivo* (32,42–44). In our study, RS67333 exhibited potent inhibition of striatal AChE, and significantly prolonged evoked striatal extracellular ACh signals, and modified activity-dependent DA release through an nAChR-dependent mechanism most likely to be desensitization.

These findings, that RS67333 inhibits endogenous AChE activity in the striatum, expand upon prior evidence that RS67333 inhibits AChE at sub-micromolar concentrations in *in vitro* assays (31). The ability of RS67333 to inhibit AChE is likely to be structurally determined. RS67333 shares key pharmacophores with donepezil, an established AChE inhibitor, including a tertiary amine that engages the catalytic anionic site (CAS) of AChE, and aromatic moieties that interact with the active-site gorge via π–π stacking interactions (31). These features have been functionally exploited in the design of donecopride, a compound incorporating structural elements of both RS67333 and donepezil, which retains dual activity as a 5-HT_4_R agonist and AChE inhibitor and has been proposed as a therapeutic candidate for treating the symptoms of Alzheimer’s disease (31). Several other 5-HT_4_R agonists, such as RS67506, prucalopride, ML10302, CJ033466, have also been reported to weakly inhibit AChE, likely owing to similar structural characteristics (31).

These findings carry several implications. Given their dual actions on 5-HT_4_Rs and AChE, RS67333 and related compounds, the effect of these compounds in brain circuits should not be considered as solely due to 5-HT_4_Rs. Some reported impacts of RS67333 on mood, memory and synaptic plasticity (7,49–51) might potentially involve AChE inhibition and not (solely) 5-HT_4_Rs. Furthermore, the identification of dual actions on 5-HT_4_ receptors and AChE also offers therapeutic opportunities. The ability to simultaneously modulate serotonergic and/or cholinergic transmission could have therapeutic potential in a range of neuropsychiatric disorders, such as in AD, PD, and depression in which these dual mechanisms may contribute helpful effects on cognition and mood.

### Alternate 5-HT_4_R ligand BIMU8 does not inhibit AChE, or alter striatal ACh or DA release

To investigate whether the effects of RS67333 seen here on striatal ACh release (and in turn DA release) are solely due to an action as an AChE inhibitor, or whether agonism of 5-HT_4_Rs can also contribute to modulation of ACh, we tested the impact of an alternative 5-HT_4_R ligand, BIMU8, on AChE activity, and in turn, ACh and DA release, at a concentration of BIMU8 that should be selective for 5-HT_4_Rs. Intrastriatal infusion of BIMU8 at a high concentration (100 µM) has been shown to modulate DA release but with some off-target effects via sigma receptors (17). The EC_50_ of BIMU8 for wild type 5-HT_4_R is reported to be ∼7-18□nM (52–54), and a concentration of 1 µM is used in the literature to activate 5-HT_4_Rs with minimal off-target effects (55). First, we established that BIMU8 (1 µM) did not detectably modify striatal AChE activity (**Fig. 4A-B**; Control: V_max_=4383 mU/mm^3^, K_m_=137.1 a.u.; BIMU8: V_max_=4278 mU/mm^3^, K_m_=129.7 a.u.; Michaelis-Menten fit). Next, we tested whether BIMU8 (1 µM) could affect ACh release in DLS detected with GRAB_ACh_. BIMU8 did not alter peak dF/F (**Fig. 4C-E**; *F*_*(1,10)*_=0.1292, *p=*0.7267, RM ANOVA main effect of the drug) or the AUC of ACh transients (**Fig. 4F-G**; *F*_*(1,10)*_=0.1422, *p=*0.7140, RM ANOVA main effect of the drug), or the decay half-life of ACh transients evoked by 1p or 5p (100 Hz) (**Fig. 4H**; *F*_*(1,10)*_=0.3155, *p=*0.5867, RM ANOVA main effect of the drug). Finally, we assessed whether BIMU8 could alter DA release detected with FCV. BIMU8 failed to modify DA release evoked by either 1p or 5p (100 Hz) stimuli (**Fig. 4I-K**; *F*_*(1,8)*_=2.036, *p=*0.1915, RM ANOVA main effect of the drug), or the 5p/1p ratio (**Fig. 4L**; *t*(4)=2.596, *p=*0.0603; Student’s paired *t*-test). These findings show that a 5-HT_4_R ligand BIMU8 at a concentration that does not inhibit AChE, does not also influence ACh or DA release dynamics, suggesting that 5-HT_4_R activation alone is not sufficient to modulate ACh or DA transmission. The lack of impact of specific 5-HT_4_R agonist BIMU8 on either AChE, ACh or DA, suggests that striatal 5-HT_4_Rs have minimal impact on these aspects of striatal circuits. The exclusion of striatal 5-HT_4_R effects in the local modulation of evoked DA and ACh release in turn indicates that the striatal actions of ligand RS67333 on DA seem solely through actions as an AChE inhibitor, altering ACh dynamics and thereby modulating DA transmission indirectly via nAChRs.

**Figure 4.**
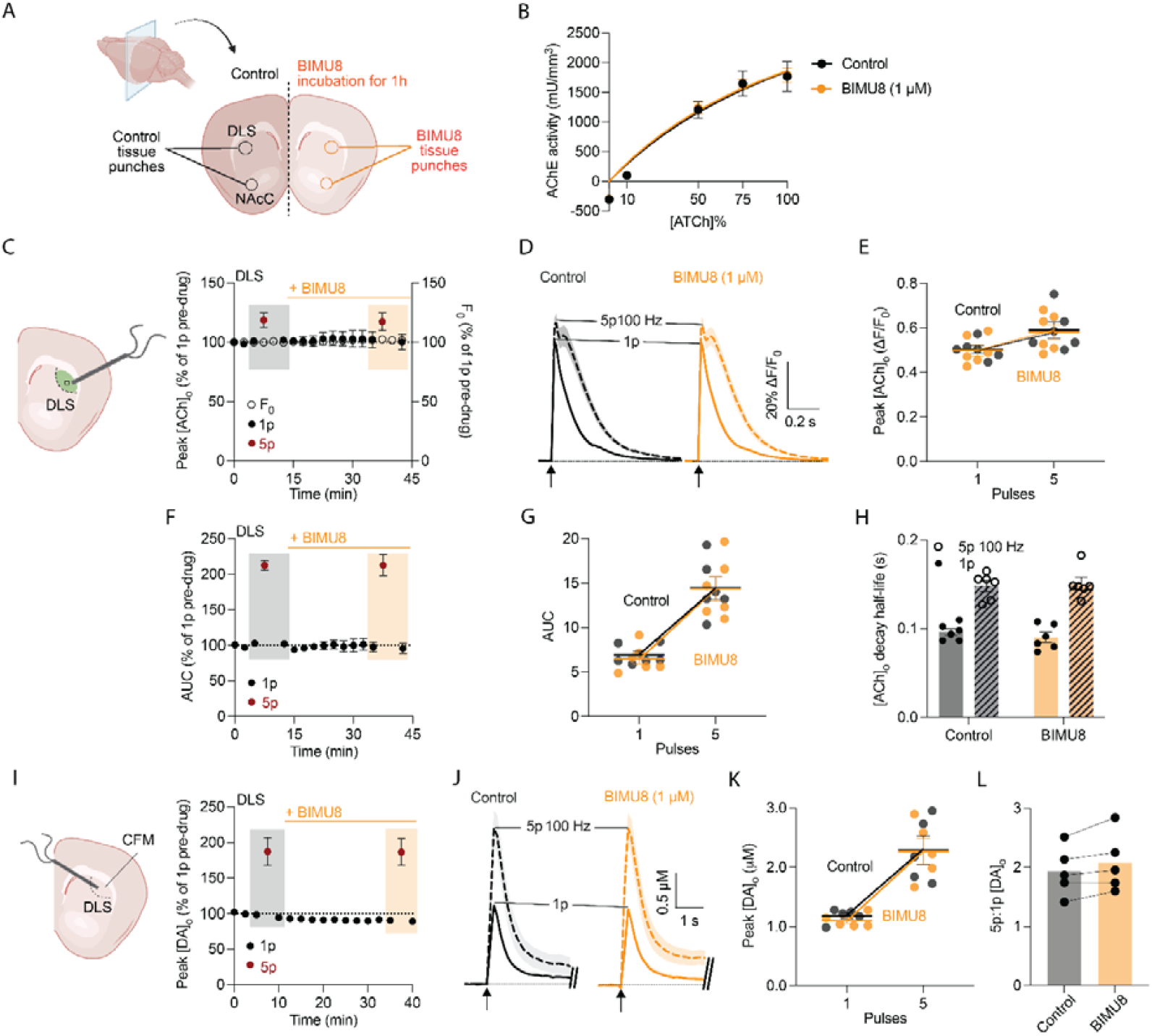
BIMU8 does not alter striatal AChE activity, or ACh or DA dynamics. **(A)** Schematic of tissue punches from striatal slices, with/without incubation in BIMU8 (1 µM) (*n*=4, 4 animals). **(B)** Fitted Michaelis-Menten curves of AChE activity. **(C)** Schematic of stimulation and GRAB_ACh_ recording, alongside normalised mean peak evoked and baseline fluorescence (F_0_) during consecutive imaging recordings of GRAB_ACh_ evoked by 1p (black) and 5p (100 Hz, red), during BIMU8 application in DLS (*n*=6, 4 mice). **(D,E)** Mean evoked ACh transients ± SEM (D), and peak ACh signals (E) before (black) and with BIMU8 (yellow) from timepoints shaded in C. **(F,G)** Corresponding mean AUCs during consecutive recordings (F) and AUC scatter plot (G). **(H)** Decay half-lives of evoked GRAB_ACh_ signals. **(I)** Schematics of stimulation and FCV recording of [DA]_o_ in DLS, alongside normalised mean peak [DA]_o_ during consecutive recordings of DA release evoked by 1p (black) and 5p (100 Hz, red), during application of BIMU8. **(J,K)** Mean [DA]_o_ transients ± SEM evoked (at arrows) (J) and scatter plot (K) for single pulses (1p, solid) and 5 pulses (100 Hz) (dashed), before (black) and with BIMU8 (yellow) (n=5, 4 mice), derived from timepoints shaded in I. **(L)** Ratios of peak [DA]_o_ evoked by 5p:1p in DLS. Schematics created in BioRender. Cragg, S. (2026) https://BioRender.com/l2hg9ad

In summary, our studies have uncovered a pronounced function of RS67333 as an inhibitor of striatal AChE that modulates striatal ACh signalling, including a downstream outcome on DA release. By contrast, striatal 5-HT_4_Rs seem to have minimal role in regulating striatal ACh or DA signalling. The potential impact of other similar purported 5-HT_4_R ligands on ACh signalling should continue to be appreciated, to prevent future misattribution of drug effects to 5-HT_4_Rs and to more fully comprehend the potential utility of such compounds. Their therapeutic utility need not become restricted; on the contrary their ‘off-target effects’ could provide therapeutic opportunities in neurodegenerative and neuropsychiatric disorders where cholinergic and dopaminergic dysfunction are involved.

## METHODS

### Animals, surgery and tissue preparation

All procedures were performed in accordance with the UK Animals (Scientific Procedures) Act 1986, with ethical approval from an Animal Welfare and Ethical Review Body at the University of Oxford, under the authority of a Project Licence (P9371BF54) from the UK Home Office. Male and female wild type C57BL/6J mice were group-housed and maintained on a 12 h light/dark cycle (lights on 07:00-19:00) with *ad libitum* access to food and water.

To prepare animals for intracranial delivery of viral vectors for the GRAB sensors for ACh or DA, mice (6-8 weeks, Charles River) were anesthetized with 4% isoflurane (O_2_ flow of ∼1 L/min) and subsequently maintained with ∼1.5-2% isoflurane during surgery. Mice were placed in a small animal stereotaxic frame (David Kopf 124 Instruments), and body temperature was maintained at ∼37°C. After exposing the skull under aseptic techniques, a small burr hole was drilled and an adeno-associated virus (AAV) solution encoding either the GRAB sensor for ACh, GRAB_ACh3.0_ (AAV2/5-hsyn-ACh_3.0,_ ≥2.00 × 10^12^ viral genome copies per ml), or for DA, GRAB_DA3h_ (AAV2/5-hysn-DA_3h,_ ≥2.00 × 10^12^ genome copies per ml) (BrainVTA, China), was injected into dorsolateral striatum (DLS) (AP +0.65 mm, ML ± 2.2 mm from bregma, DV −2.7 to -2.5 mm from exposed dura mater) and/or nucleus accumbens core (NAcC) (AP +1.4 mm, ML ± 0.9 mm from bregma, DV −3.8 to -3.5 mm from exposed dura mater) (1 μl per hemisphere) at an infusion rate of 200 nl/min, using a 32-gauge Hamilton syringe and needle (Hamilton Company). The needle was left *in situ* for 5 minutes post injection. Animals were allowed to recover and were culled by cervical dislocation for fluorescence imaging 3-4 weeks post-surgery.

For *ex vivo* recordings, male and female C57BL/6J mice (8-18 weeks, Charles River) underwent cervical dislocation and decapitation and the brains removed and transferred to an ice-cold cutting solution, containing in mM: 194 sucrose, 30 NaCl, 4.5 KCl, 1 MgCl_2_-6H_2_O, 26 NaHCO_3_, 1.2 NaH_2_PO_4_, 10 glucose, saturated with 95% O_2_/5% CO_2_. Coronal slices 300 μm-thick containing the striatum were cut using a vibratome (VT1200S, Leica Microsystems). Slices were immediately transferred to artificial cerebrospinal fluid (aCSF) containing in mM: 130 NaCl, 2.5 KCl, 26 NaHCO_3_, 1.25 NaH_2_PO_4_, 2 MgCl_2_-6H_2_O, 2.5 CaCl_2_ and 10 glucose, saturated with 95% O_2_/5% CO_2_. Sections were incubated at 34°C for 15 min before they were stored at room temperature (20-22°C) until recordings were performed. All recordings were obtained within 6 h of slicing. Both sexes were included and balanced throughout experiments.

### Fast-scan cyclic voltammetry

Evoked extracellular DA concentration ([DA]_o_) was measured as previously (28) in acute coronal brain slices using fast-scan cyclic voltammetry (FCV) at carbon fibre microelectrodes (CFM) fabricated in-house. In brief, acute striatal brain slices were hemisected and transferred into a recording chamber and superfused at ∼2.5 ml/min with aCSF at 31-33°C. A CFM (diameter 7-10 μm, tip length 50-150 μm) was inserted 100 μm into the tissue and left to equilibrate for 50 min prior to recordings. All experiments were carried out either in the dorsolateral quarter of dorsal striatum (dorsolateral striatum, DLS) or in nucleus accumbens core (NAcC) (within 100 μm of the anterior commissure), one site per slice. The sampling of these two regions allows sampling spanning regions that represent functional extremities of mid-anterior striatum. For each experiment, data were collected from at least 3 animals and 5 slices. [DA]_o_ evoked by local electrical stimulation using a concentric bipolar Pt/Ir stimulating electrode (FHC Inc.) were measured using FCV with a Millar voltammeter at CFMs. A triangular voltage waveform (-0.7 to +1.3 V vs. Ag/AgCl, scan rate 800 V/s) was applied, with a sampling frequency of 8 Hz. Electrical stimulation (0.6 mA, 0.2 ms pulse width) was delivered every 2.5 minutes (DS3 constant current isolated stimulator, Digitimer). Evoked currents were confirmed as DA due to their characteristic oxidation (500 – 600 mV) and reduction (-200 mV) potentials. All FCV data were acquired using Axoscope 10.6 (Molecular Devices) and analysed using local written Python script. Electrodes were calibrated at the end of the experimental day in 2 μM DA, prepared immediately beforehand in recording solutions. DA oxidation currents values were measured from background-subtracted voltammograms, converted to concentration using the electrode calibration factor. The drug RS67333 (but not BIMU8) was found to reduce the sensitivity of the CFM to applied DA to 54 ± 2 % of the electrode sensitivity calibrated in the absence of RS6733 (n=29 electrodes). [DA]_o_ recorded in the presence of RS67333 were therefore scaled using a standardised coefficient of 100/54.

### GRAB imaging

GRAB sensors were imaged as described previously (56). In brief, acute striatal brain slices were hemisected and transferred into a slice immersion recording chamber and superfused at ∼2.5 ml/min with aCSF at 31-33°C. A 10X water immersion objective (Olympus) was lowered into the superfusate and an LED (470 nm, 4 mW, pE-300, CoolLED) was briefly switched on to visualise GFP expression to confirm expression, and to place the stimulating electrode. All images were recorded by a microscope (SliceScope, Scientifica) and a Prime BSI Express Scientific CMOS camera (Teledyne Photometrics). Sampling rate was 100 Hz, exposure time 10 ms, field size was 665.6 μm × 665.6 μm and images sequences were recorded for 5 s with a 1 s pre-recording photo-bleaching time using Micro-Manager2.0. Electrical stimulations, LED light, and image acquisition were synchronised using TTL-driven stimuli via Multi Channel Stimulus II (Multi Channel Systems). Electrical stimulations were delivered every 2.5 minutes. Fluorescence of GRAB_ACh_ or GRAB_DA_ was captured from the region of interest (ROI, 50 µm × 50 µm), at a distance of 50 µm from the stimulating electrode tip. Data are expressed as a change in fluorescence from baseline (ΔF/F_0_), where F_0_ is the baseline signal prior to the stimulation. To evoke ACh or DA release, electrical stimulus pulses (single pulses, 1p, or 5 pulses at 100 Hz; pulse duration, 200 μs; 0.6 mA) were given by a local bipolar concentric Pt/Ir electrode. Image files were analyse with MATLAB and ImageJ.

### AChE assay

Slices of 300 μm-thick acute coronal striatal slices were prepared using a vibratome in ice-cold cutting solution as described above. Slices were bisected, and opposite hemispheres incubated in either aCSF control, 10 µM RS67333 or 10 µM BIMU8 for 1 hour at room temperature. Dorsal and ventral striatal tissue punches (1.2 mm diameter) were collected and AChE activity was measured using a commercially available colorimetric assay kit (Abcam, ab138871) which utilizes 5,5′ -dithiobis-(2-nitrobenzoic acid) (DTNB) to detect thiocholine (TCh) produced from acetylthiocholine (ATCh) hydrolysis by AChE. Tissue punches were lysed in PBS containing 0.5% IGEPAL CA-630 for 10 minutes, sonicated (10-15 s, 2-3 cycles), and centrifuged at 800 rpm for 5 minutes at 4□°C. Supernatants were diluted 1:5 in assay buffer. ATCh-containing reaction mixtures were loaded into wells of a 96-well plate following addition of 50 µL of diluted sample and incubated for 15-minutes incubation at room temperature in the dark. Absorbance was measured at 410 nm using a microplate reader (PHERAstar FSX). Saturation curves were generated based on the calculated enzyme activities. The test compounds (RS67333 or BIMU8) were present in the reaction buffer throughout the entire assay process. Michaelis constant (K_m_) and maximum reaction velocity (V_max_) were determined by fitting the data to the Michaelis–Menten equation using non-linear regression.

### Drugs

RS67333 was obtained from MedChemExpress (MCE), dihydro-β-erythroidine hydrobromide (DHβE) and BIMU8 were obtained from Tocris Bioscience. Drugs used in acute recordings were diluted to their required concentrations in aCSF immediately before use. RS67333 is typically used at micromolar concentrations in ex vivo slices to assess 5-HT_4_R function (57,58), and thus concentrations of 1-10 µM were used here in mouse brain slices. *In vitro*, RS67333 inhibits human AChE with an IC_50_ of ∼403 ± 86□nM (31).

## Data analysis and statistics

FCV data were acquired using Axoscope 10.5 (Molecular Devices) and analysed using locally written Python code. GRAB data were acquired and analysed using Image J and local written MATLAB scripts. Statistical analysis was carried out using GraphPad Prism v7. Data sets were analysed using Shapiro-Wilk test for normality. No outlier data were identified or removed. Data sets with normal distributions were analysed for significance using paired Student’s two-tailed *t*-test or analysis of variance (ANOVA) measures followed by Fishers LSD post-*hoc* test. Data are expressed as mean ± standard error (SEM). n indicate number of experiments=slices (in number of mice). The number of animals for each dataset is N≥3.

## Data availability

The source data, as well as protocols, key lab materials, and code used and generated in this study are listed in a Key Resource Table and available alongside their persistent identifiers at http://doi.org/10.5281/zenodo.17278002.

## Acknowledgements

We thank Dr Katherine Brimblecombe for support with FCV, Prof. Richard Wade-Martins and Dr Kaitlyn Cramb for access to a microplate reader, and the staff in Biomedical Services for veterinary support. We thank Dr Yizhou Zhuo for helpful discussions about the use of GRAB_DA_.

## Author Contributions

QQ and SJC designed the experiments, QQ and WW performed experiments, QQ analysed data, QQ and SJC interpreted the results, wrote and revised the manuscript.

## Funding

The work is supported by Aligning Science Across Parkinson’s (ASAP) through the Michael J. Fox Foundation for Parkinson’s Research (ASAP-020370, ASAP-025192).

## Competing Interests

QQ and WW declare that the research was conducted in the absence of any commercial or financial relationships that could be construed as a potential conflict of interest. SJC is a consultant to Oxneuro.

## Notes

### Competing Interest Statement

The authors have declared no competing interest.

